# Comparing random and timed mating of mice in the experimental paradigm

**DOI:** 10.1101/2023.12.16.571983

**Authors:** M Imran, M Mohsin, T Azam, K Fatima, MK Naeem, W Ahmad, R Basri, S Naz, M Anwar, Q Jabeen, M. Khurshid

## Abstract

*Mus Musculus* has provided scientists a well-adapted animal model for the developmental studies. There is one on one co-relation between molecular determinants such as genes between humans and mice hence making them superior model to study gene expression and to do translation research. Swiss albino mice strain of house mice was used in this study with comparisons of efficiency of pregnancies during two different strategies of mating; i.e., more traditional random/blind mating and more recent timed mating in experimental paradigms to get staged mice embryos. The workflow required for the blind mating was less labor intensive and required minimal maintenance both time and human resource and was easy to execute. On the other hand, timed mating strategy was ideal for getting staged embryos that was efficient in giving pregnancies and was ideal for less equipped labs but however required more maintenance and human resource. The results of comparisons of weight gain with pregnancies was similar in two groups where some un-pregnant mice also showed average or above average weight gain with time. The z-test analysis for difference between two proportions was employed to calculate efficiency of pregnancies in the two groups and p value was found to be 0.06 which suggested that was no significant difference between the two groups however, this might be due to small sample size that was possible in our small setting. The accurate embryonic staging was required in the experiments corresponding to the peak of neurogenesis in mice and embryos were fixed and sectioned and later studied with immunohistochemistry. The timed matings showed to be superior regarding total pregnancies and having accurate stages and is recommended in labs even if in smaller settings and prevent less animal sacrifice.

## Introduction

The scientific name of house mice is *Mus Musculus* and it belongs to kingdom Animalia, phylum chordate, class Mammalia and order Rodentia. Mice are preferred in Genetics and Molecular biology research because they show great similarity with humans with respect to primordial genetic, physiological and anatomical components (1). To investigate human health almost 59% animals used in biomedical research are mice (2). Short life cycle, short gestation period and high breeding efficiency make them an excellent animal models in laboratory settings (3). Moreover, mice are small, easy to handle and can be transported easily. They are used to study human diseases and human genetics because they are phylogenetically related to humans (4). Gestation periods of mice is between 18 to 22 days (5). The developmental biology studies require accurate staging of embryos that can help with comparative developmental studies between mice with mice or mice with different species known as evo-devo approach (6). Almost 95% of the genes of human and mice are similar so it is an excellent model to do gene expression studies involving transcriptional analysis (7). To get staged embryos mice may be mated randomly or at specific stages of estrous cycle. We anticipated to compare both strategies in smaller settings with respect to efficiency of pregnancies and accurately staged embryos for downward analysis. It looked apparent that the mating on the day of ovulation; i.e., at the proestrous-estrous and visualization of plug the next day will lead to more efficient pregnancies and also accurate staging if pregnant. The estrous cycle of female mice consists of four stages namely proestrous, estrous, metestrous and diestrous (8). There are various methods to evaluate the estrous cycle of mice that include visual observation (9), vaginal cytology (10), histology of reproductive organs (8), vaginal wall impedance (11) and biochemistry of the urine (12, 13). We employed visual observations of the vagina and vaginal cytology combinations and set out on this comparative study.

## Material and Method

### Animal acquisition

The collaboration letter was signed between the Department of Biochemistry and Department of Pharmacy to do the animal work. The Islamia University of Bahawalpur’s Ethical committee approved the use of animals in the work via letter no. 3990/ORIC. We acquired albino house mice, BALB/c that were enrolled in the study. We enrolled mice n=15 blind/random matings experiment and n=15 in second experiment of timed matings in mice. Male to female ratio was 1:2.

### Housing of the animals

We set up two experiments different experiments scheduled at different time periods to get timed embryos at different embryonic days (from E12.5-E18.5). In the first experiment we took 10 female mice in a separate cage and housed 5 stud males mice each in a separate cage to increase their sperm level, fertility and also to avoid their aggressive behavior (14). We used wooden cages that were custom made and had a proper fresh water supply with adjusted bottles. The pelleted food was given which was made up of ground ingredients. We used fine wooden scrap for the cage beddings. The cages were placed in the air-conditioned environment to maintain the optimum temperatures in summers. In second experiment we used the same setting and numbers but were analysed for estrous stage.

### Experiment 1: Random or blind mating

1. Initially, female mice were labeled by using green and black permanent makers. One, two, three and four black bands on the tail of mice represent mice number 1, 2, 3 and 4 respectively while full black tail show mice number 5. Similarly one, two, three and four green bands represent mice number 6, 7, 8 and 9 respectively. Further full green tail was identified as mice number 10.
2. Body weight of all female mice was measured in the start of the experiment. Then all female mice were shifted to the male cages in1:2 male to female mice ratios. Vaginal plug was checked in next day early in the morning. The female mice in which vaginal plugs were observed were separated from the males and were separated.
3. Weight of all female mice were then recorded on the daily basis (whenever possible).
4. At the desired embryonic day after the separation the mice were dissected to collect the embryos (the day of plug was considered 0.5 embryonic day) and adult tissues.

### Experiment 2: Timed mating

1. Method of labeling the mice in this trial was same as above and the weights of each female mice were also recorded on daily basis (whenever possible).
2. To evaluate estrous stage of mice we used vaginal cytology protocol (as per Byers et al. 2012 (8) using saline dropper method and but the smear staining was done by crystal violet staining and visualized by light microscopye) combined with visual observations under light was done (8).
3. Female mice that were in their pro-estrous to estrous stages were set for mating.
4. The next day, in the early morning, the female in which vaginal plug was visible were separated and dissected at the required embryonic day to get staged embryos (the day of plug was considered 0.5 embryonic day).
5. The remaining female mice were also mated using same strategy until all mice were mated and dissected respectively.

### Dissection of mice

On the desired embryonic day the mice were dissected to collect the embryos from the pregnant dams. For downward experimentation the embryos were collected either in RNA later or fixed in formalin and some embryos were also snap frozen. We did this for future work.

### Statistical analysis

Two proportion Z-test was used to calculate the efficiency of pregnancies in the two independent groups and p values were calculated.

### Immunohistochemistry

The formalin fixed paraffin embedding and sectioning was done and later stained by Immunohistochemistry using established (15) and improved protocols. DAPI (4′,6-diamidino-2- phenylindole) a counterstain that binds to A/T rich DNA bases in the grooves is used where it marks the complete architecture and cellular organisation in the tissues. Embryonic tissues at various stages were stained.

## Results

### Weight dynamics and pregnancies

Weight gain is one of the hallmarks of pregnancy generally but in a small animal like mice it sometimes is not relevant. Here, we analysed weight dynamics in both the experiments oin the first 12 days in mice (the starting embryonic day at which embryos were anticipated to be collected) and was found to be not related to the induction of pregnancy. The weight readings considerably increase in pregnant mice after day 12 of pregnancy and would give false indication if weight dynamics were matched of E12.5 and E18.5 mice hence the period was kept constant. Female mice were also plotted for total weight gain calculated from the initial weight at the start of trial and before dissection and was plotted (see figure 1). Mice in both groups showed weight gain ranging from 4g to 18g but found to be not co-related directly with pregnancy (see tables 1, 2 and 3). However, mice with higher than 6g of net weight gain corresponded well with pregnancies.

**Figure 1:**
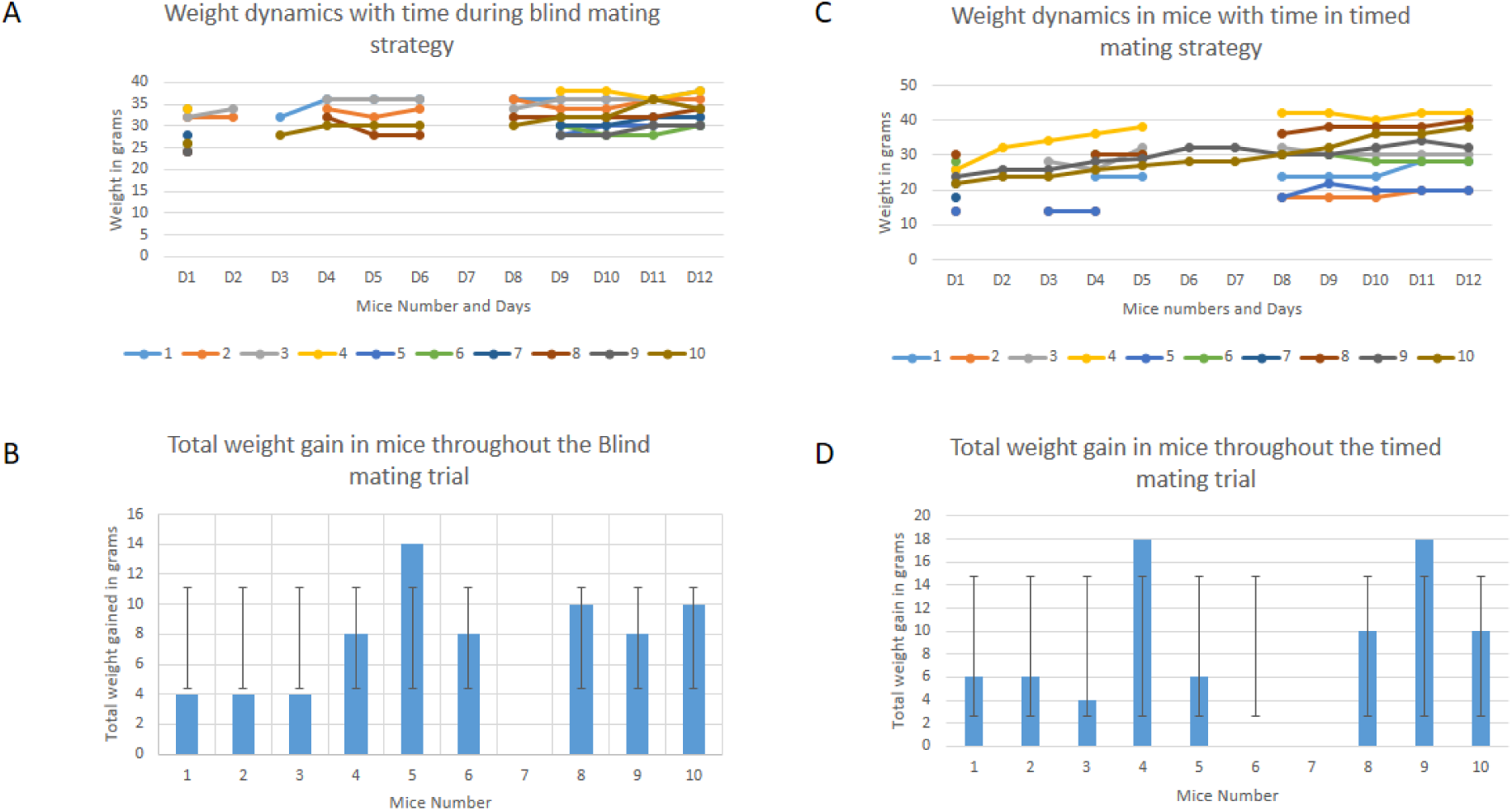
A, Weight dynamics in female mice during blind mating trial from Day1 to Day12. **B,** Total weight gain in mice throughout the blind mating trial. **C,** Weight dynamics in female mice in timed mating trial from Day1 to Day12. **D,** and Total weight gain in mice in timed mating trial

**Table 1.** Correlation of mice with weight during the two experiments.

| Mice No | Initial weight (g) |  | Final weight before dissection (g) |  | Net weight gain (g) |  |
| --- | --- | --- | --- | --- | --- | --- |
|  | Exp1 | Exp2 | Exp1 | Exp2 | Exp1 | Exp2 |
| 1 | 34 | 22 | 38 | 28 | 4 | 6 |
| 2 | 32 | 14 | 36 | 20 | 4 | 6 |
| 3 | 32 | 26 | 36 | 30 | 4 | 4 |
| 4 | 34 | 26 | 42 | 42 | 8 | 18 |
| 5 | 24 | 14 | 38 | 20 | 14 | 6 |
| 6 | 26 | 28 | 34 | 28 | 8 | 0 |
| 7 | Paralyzed | Died |  |  |  |  |
| 8 | 24 | 30 | 34 | 40 | 10 | 10 |
| 9 | 24 | 24 | 32 | 32 | 8 | 8 |
| 10 | 26 | 22 | 36 | 40 | 10 | 18 |

**Table 2:**
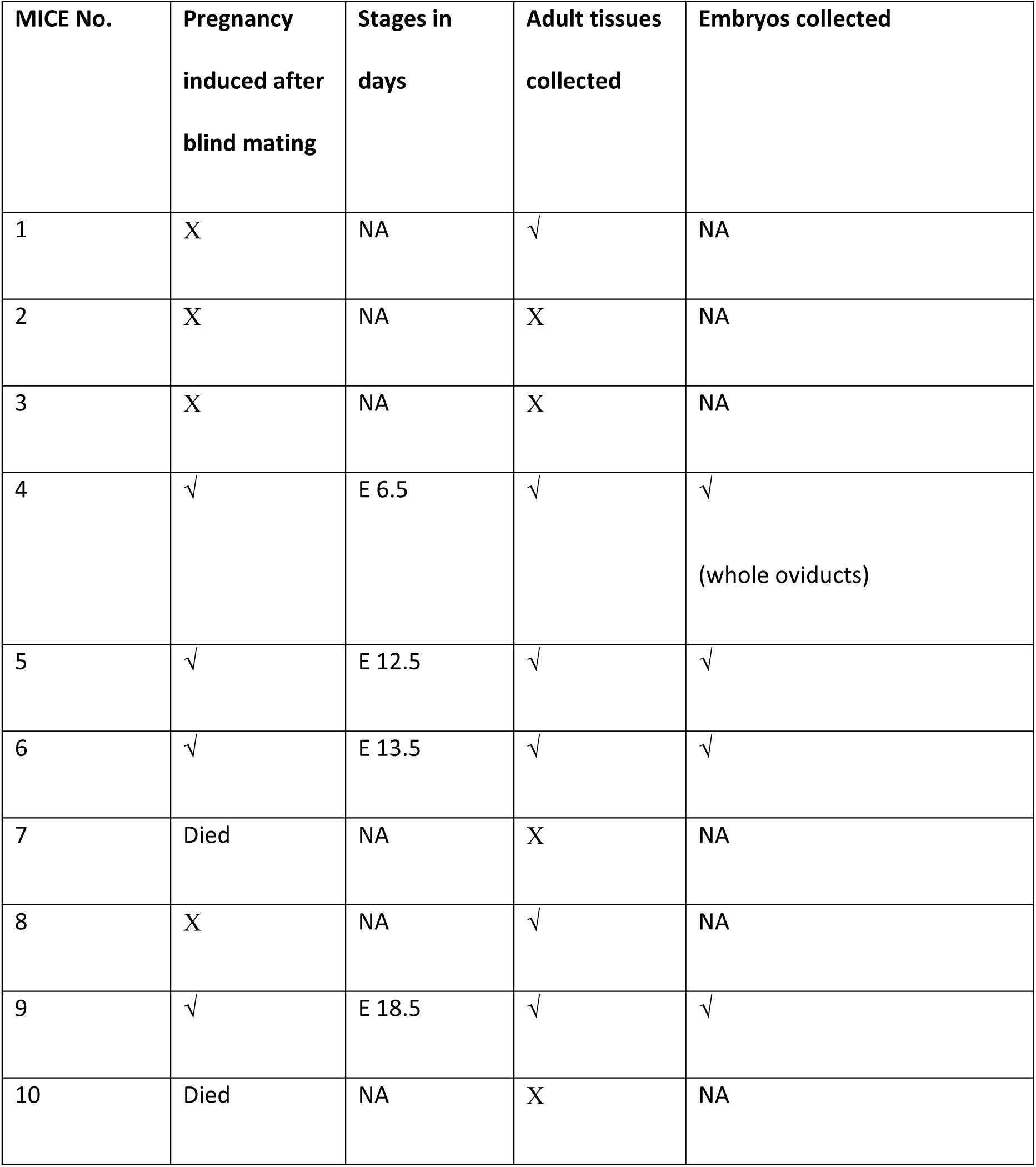
Random or blind matings and the success in pregnancies and collection of samples.

| MICE No. | Pregnancy induced after blind mating | Stages in days | Adult tissues collected | Embryos collected |
| --- | --- | --- | --- | --- |
| 1 | X | NA | √ | NA |
| 2 | X | NA | X | NA |
| 3 | X | NA | X | NA |
| 4 | √ | E 6.5 | √ | √<br>(whole oviducts) |
| 5 | √ | E 12.5 | √ | √ |
| 6 | √ | E 13.5 | √ | √ |
| 7 | Died | NA | X | NA |
| 8 | X | NA | √ | NA |
| 9 | √ | E 18.5 | √ | √ |
| 10 | Died | NA | X | NA |

**Table 3:** Timed pregnant mice results showing more pregnancies and accurate embryonic stages.

| MICE No. | Pregnancy induced after timed mating | Stages in days | Adult tissues collected | Embryos collected |
| --- | --- | --- | --- | --- |
| 1 | √ | E 14.5 | √ | √ |
| 2 | √ | E 7.5 | √ | √<br>(Whole embryos) |
| 3 | √ | E 13.5 | √ | √ |
| 4 | √ | E 14.5 | √ | √ |
| 5 | √ | E 15.5 | √ | √ |
| 6 | √ | E 15.5 | √ | √ |
| 7 | Died | NA |  |  |
| 8 | √ | E 13.5 | √ | √ |
| 9 | √ | E 13.5 | X | √ |
| 10 | X | NA | X | X |

Weight gain was observed in each mice, though it did not reflected true pregnancy (Table 1).

### Blind mating trial and number of pregnancies

(Blind mating experiment of 10 female mice with 5 stud male mice took place from 16-05-22 to 20-06- 22) (see table 2). The vaginal plug was observed once and female mice were separated but it led to false pregnancies or/and non-pregnant mice in more than half of female mice that were sacrificed. In total we got 40% pregnant rate (see table 2).

### Timed mating trial setup and number of pregnancies

(Timed mating experiment took place in two phases from 24-03-23 to 10-05-23 and later in 2024). As above total of 10 female mice were mated with 5 stud male mice (Table 3) Visual observations of the vagina in female mice are different at different estrous cycle stages and hence maybe used to assess the cycle stage (Figure 2A). In pro-estrous stage the vaginal tissues and surrounding tissues appear pink, moist and swollen with a wide opening and with wrinkles in dorsal and ventral edges whereas, at estrous stage the vaginal tissues are relatively less swollen and are not pink or moist and are ideal for mating. However, during metestrous stage, the vaginal opening is small and has minor swelling and at diestrous no opening and swelling of vagina was observed (Figure 2A). In addition, the vaginal cytology of mice at various estrous cycle stages may be done to determine the estrous cycle stages (Figure 2B). The vaginal cytology at pro-estrous stage showed majority of nucleated cells and small numbers of cornified cells whereas the estrous stage was revealed by large numbers of cornified epithelial cells. The metestrous was marked with cornified epithelial cells plus polymorphonuclear leukocytes with some nucleated epithelial cells and in diestrous, which lasts for 2 days and is the longest stage in estrous cycle there are more polymorphonuclear leukocytes (see figure 2B). The combination of both the observations was done to identify female mice at pro-estrus to estrous stage that lasts up to 15 hours and represent the day of ovulation and were set to mating with stud males (see supplemental data). The female mice with vaginal plug were separated next day to get high efficiency of pregnancy and accurately staged embryos. The number of pregnancies as expected were quite high as anticipated and was 80% in this group (see table 3).

**Figure 2:**
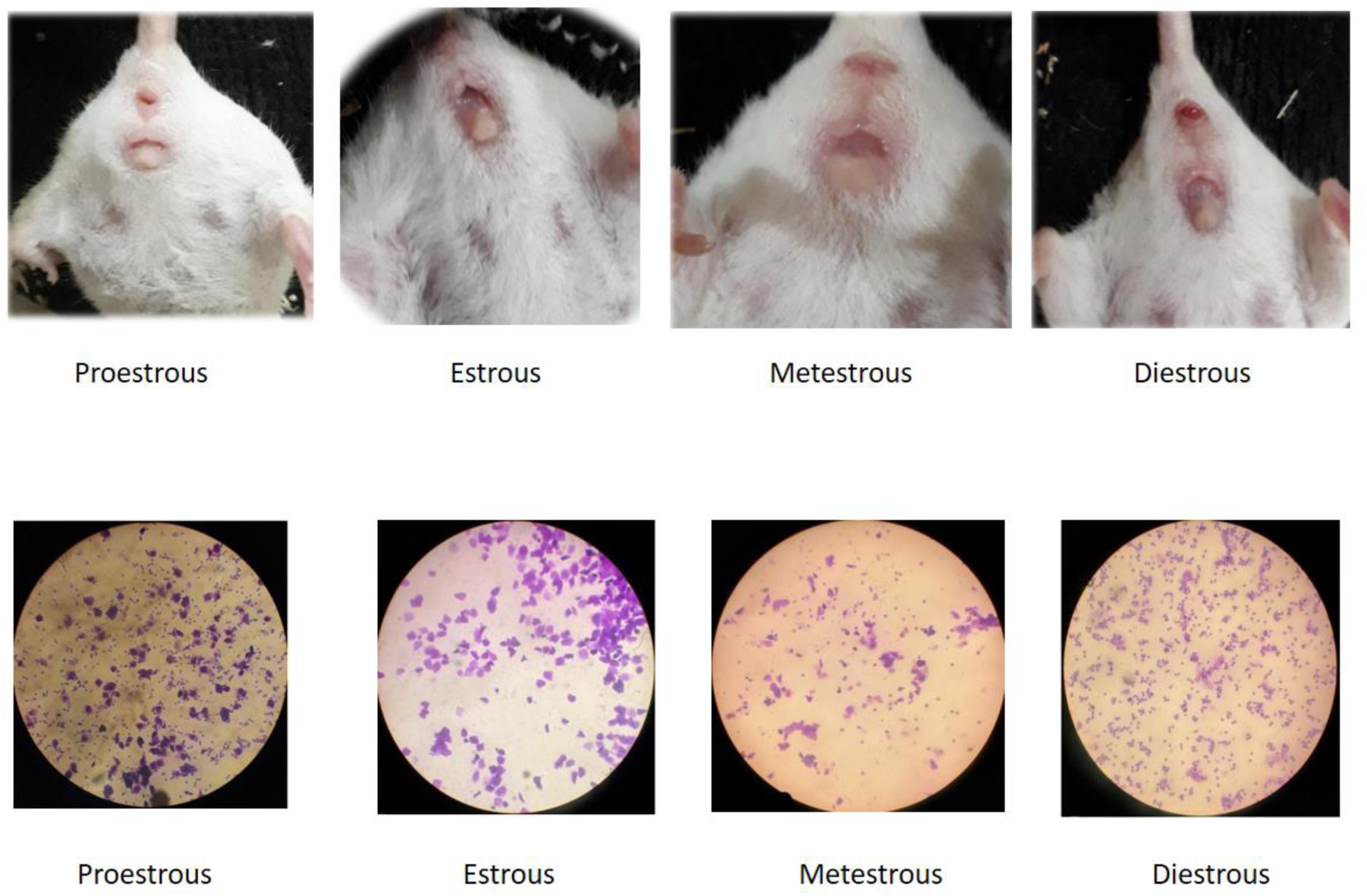
A, The shape of vagina changes at various stages in the mice estrous cycle through visual observations, **B,** The stages are visible after crystal violet dye staining of the vaginal smear at various stages in the mice estrous cycle

### Two proportion z- test analysis to calculate statistical difference if any in rate of pregnancies in both groups

A two proportion z- test is used to calculate the true difference between two independent groups and to a given confidence interval. We used this analysis to calculate statistical difference in the rate of pregnancies in both groups. Our data was shown to have p value of 0.06 and correspond with the null hypothesis which states both groups are same hence there is no statistical difference in these two mating strategies in terms of efficiency of pregnancy in both groups (Figure 3). This may be due to small sample size in the two groups. This was unavoidable due to small laboratory setting and availability of mice.

**Figure 3:**
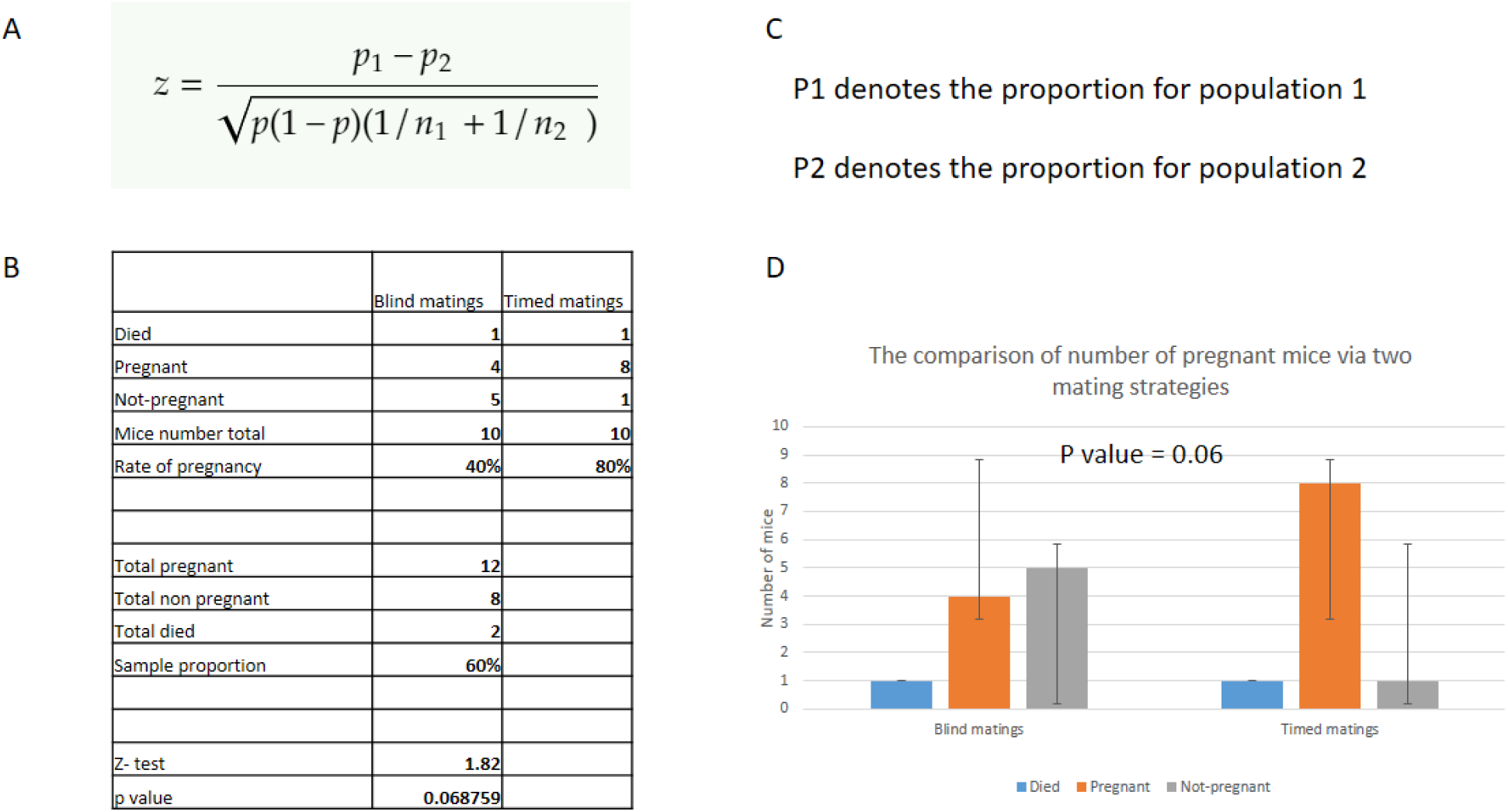
**A**, z test to calculate p values for two proportions analysis. **B-C,** P values calculated for number of pregnancies between blind and time mating was done. **D,** 0.06 p value between two groups support null hypothesis is correct.

### Embryonic Stages collected and downwards analysis

Embryonic stages were collected and were verified by characteristic feature of embryos at respective stages (see figure 4).

**Figure 4:**
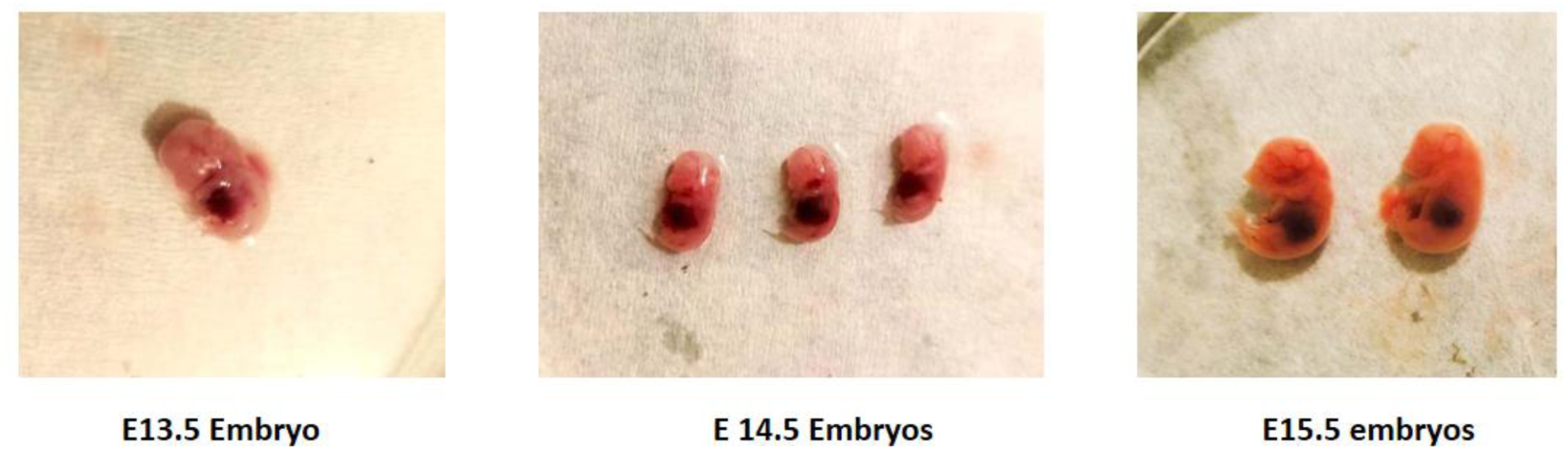
The embryo stages collected during the experiment E13.5, E14.5 and E15.5 (Left to right.

The embryo at 13.5 stage (see Figure 4) has foot plate that is anterior and has posterior distal borders that are sectioned. Hand plate has more clearly sectioned than foot plate. In hind limb only ankle region is identifiable while in forelimb elbow and wrist are distinguishable. Furthermore pinna and whiskers are appearing.

At E14.5 embryonic stage (see Figure 3) the separation of individual digits in the anterior footplate is a unique feature of this stage. Although there isn’t yet any discernible separation between the growing toes and the rear footplate exhibits significant indentations.

At embryonic stage E15.5 (see Figure 4) there was complete interdigital indentation is the hand and foot plates, and the vanishing of the webbing improves the delineation of each individual digit. The elbow and knee are obviously bent. Hair follicles can be found in the cephalic region, but not along the margins of the vibrissae. The anterior area of the back is gray, and the curvature of the back is markedly different from the currently straight post-cranial vertebral axis.

### Immunohistochemistry to confirm the staging of embryos

Downwards analysis for confirmation of embryonic stages was done using formalin fixation and paraffin embedding followed by sectioning and immunohistochemistry. DAPI (4′,6-diamidino-2-phenylindole) staining was used to assess the stage by developing neuroepithelium which shows overall architecture of all cell nuclei. The figure 5 shows that E12.5, E13.5 and E14.5 embryos showing developing neuroepithelium where the formation and organization of layers confirmed the stage.

**Figure 5:**
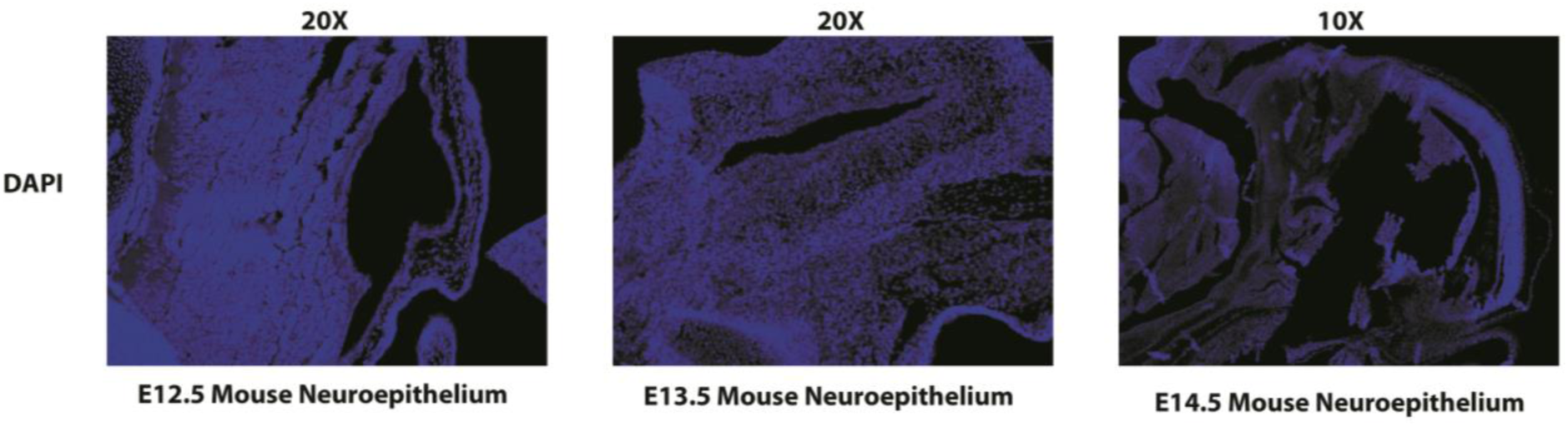
Immunohistochemistry of paraffin embedded sections of mice embryos showing developing neuroepithelium at the peak of neurogenesis is shown that were acquired through the timed mating experiment.

## Discussion

In the series of experiment done in this study we got contradicting results showing no/very little dependency on weight gain with the successful pregnancies. Average net weight gain in female mice in both random and blind matings was 4g regardless of pregnancy induction. However in the pregnant mice females with net weight gain of higher than 6g correlated with higher proportions of pregnancies in both groups. The two trials were done at different times but in the same settings. The pregnancy number was two folds in timed matings as compared to random/blind mating which was evident from mice behaviour and other features in addition to weight of female mice but not only weight gain. This was due to the mating exactly at the day of ovulation which is at pro-estrous/estrous stages and increases chances of pregnancy. In blind matings we were only sure of staging after dissecting out embryos and didn’t always find the required stage which maybe due to dropping out of vaginal plug and mating again however the timed mating experiment took more time and was labour intensive corresponded well with efficient pregnancies. The values of weight gains in both experiments were almost comparable but just on the criterion of weight it is hard to judge the pregnancy and more so the exact stage we are looking for as was our requirement in our study and hence were dependent on the observation of vaginal plug. In one study the early pregnancy around E7.5 was found to be more related to significant weight gain compared with non-pregnant mice (16) our data doesn’t support it. The time taken to finish the first experiment was 4 weeks whereas timed mating took almost two months to complete due to staging confirmation prior to sacrifice and required more housekeeping of mice and care for the animal requirements but inturn gave effcient pregnancies and staging which was confirmed by downwards immunohistochemistry analysis using DAPI stain.

The second question was whether we can statistically calculate difference between the proportions of pregnancies in the two groups. For this we used, Two proportions Z-test and although looking quite different in numbers the p value came out to be 0.06 which is not statistically significant and supports the null hypothesis which states both groups have same rate of pregnancies. This may be due to small sample size which was unevitable due to small setup and was unavoidable and might not reflect the true difference. In previous work by Byers et al. (2012) and others (17,18) with more than 100 sample size and mating variations and and some small group mating trials, there was some statistical difference in the two groups which seem to be the true reflection of the strategies in large cohorts. Our numbers also reflect this but states are different because of very small sample size. In bigger lab settings, with high resolution microscopy random/blind mating might be continued to be used as traditionally used where staging of embryos maybe confirmed by developmental hallmarks but in small setting, timed mating is relatively a method of choice for staged embryos and efficient pregnancies and might be adopted.

## Conclusions

The timed matings gave us more pregnant mice at desired stages than mice with blind matings in terms of numbers however statistically there was no significant difference between two groups and supports null hypothesis. However, it might still be adopted in smaller setting to get efficeient and accurate staging of embryos.

## Acknowledgements

We thank out whole research team and HEC, Pakistan for providing funds for this project. We especially thank Professor Qaisar Jabeen, Department of Pharmacy for allowing us to work in the animal house facility. Furthermore, microscopy was done at Shifa international hospital Islamabad and we want to thank the team there.

## Supplementary Data

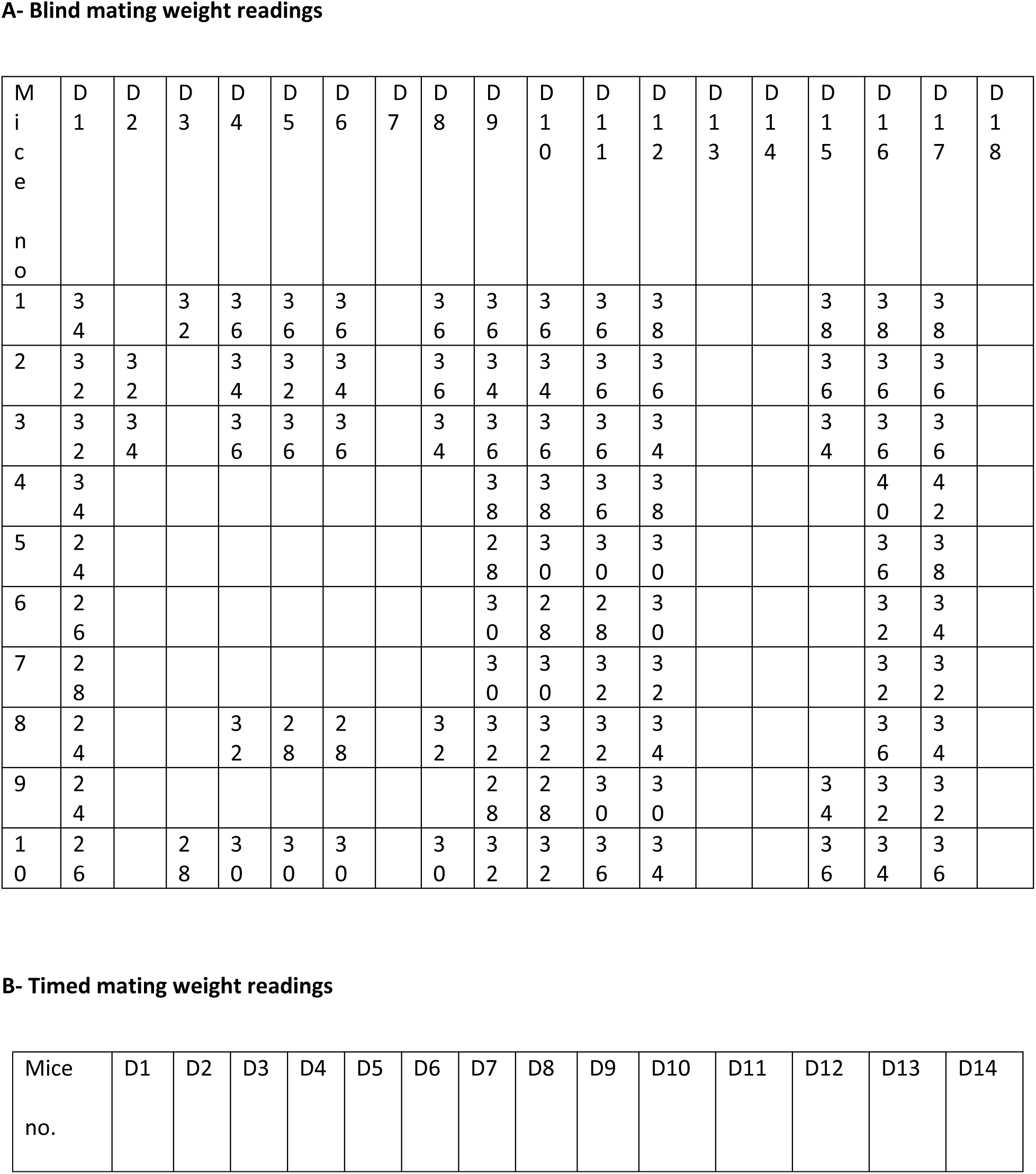
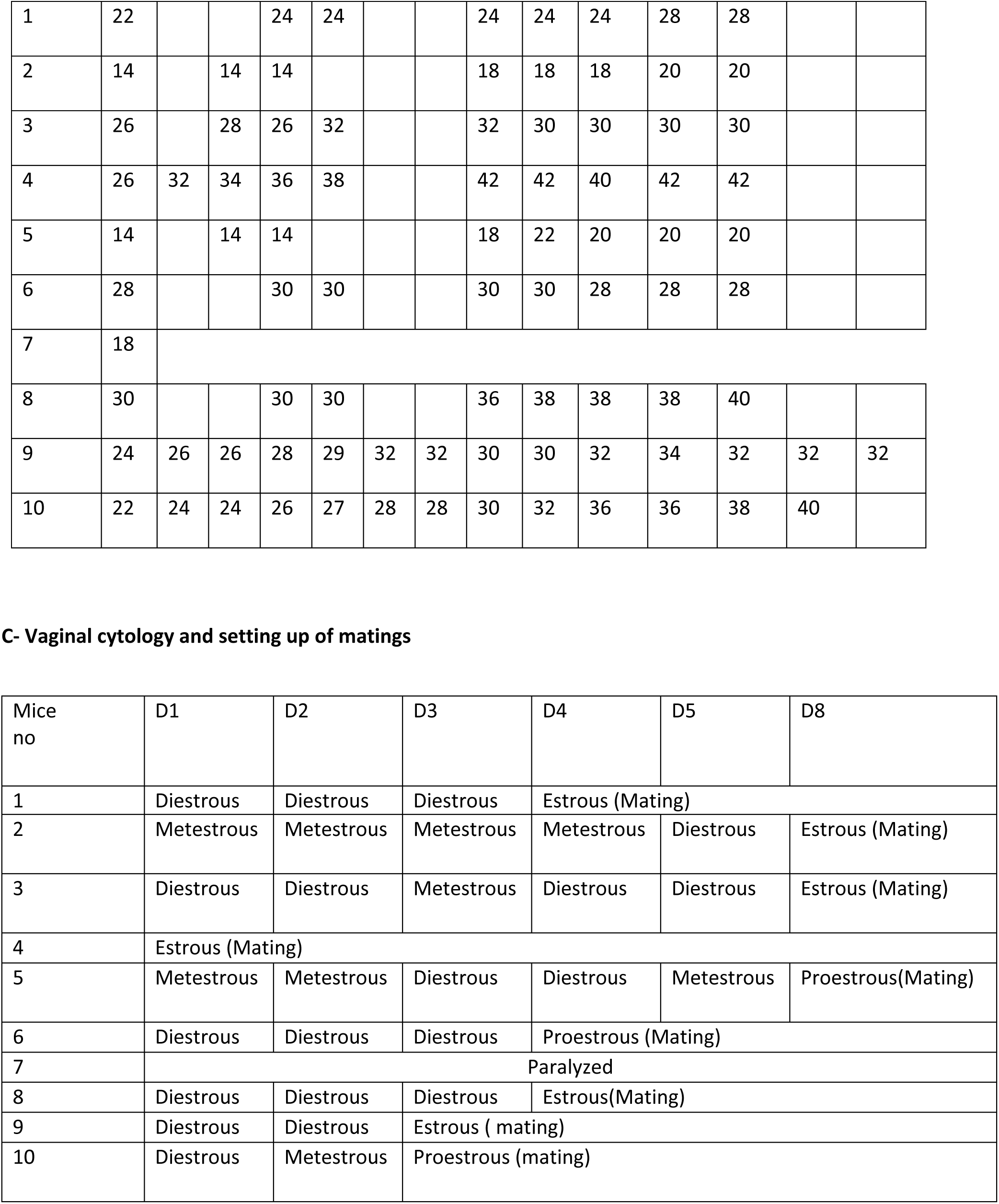

## Notes

### Competing Interest Statement

The authors have declared no competing interest.

### Summary of Updates

The manuscript was updated by adding the detailed analysis of efficiency of pregnancies in both independent groups which was not possible before due to the unequal sample size in the two groups due to constraints in the availability of mice in the season and other issues. Furthermore, the statistical test was also added after the sample sizes have become equal. Downward analysis of the embryonic tissues not possible before is also added. Some more authors have put in their contributions and are hence added.

