## Supplementary table1,2 and 3 for "Comparing random and timed mating of mice in the experimental paradigm"

**Supplementary Data:**

**A- Blind mating weight readings**

| Mice no | D1 | D2 | D3 | D4 | D5 | D6 | D7 | D8 | D9 | D10 | D11 | D12 | D13 | D14 | D15 | D16 | D17 | D18 |
|---|---|---|---|---|---|---|---|---|---|---|---|---|---|---|---|---|---|---|
| 1 | 34 | | 32 | 36 | 36 | 36 | | 36 | 36 | 36 | 36 | 38 | | | 38 | 38 | 38 | |
| 2 | 32 | 32 | | 34 | 32 | 34 | | 36 | 34 | 34 | 36 | 36 | | | 36 | 36 | 36 | |
| 3 | 32 | 34 | | 36 | 36 | 36 | | 34 | 36 | 36 | 36 | 34 | | | 34 | 36 | 36 | |
| 4 | 34 | | | | | | | | 38 | 38 | 36 | 38 | | | | 40 | 42 | |
| 5 | 24 | | | | | | | | 28 | 30 | 30 | 30 | | | | 36 | 38 | |
| 6 | 26 | | | | | | | | 30 | 28 | 28 | 30 | | | | 32 | 34 | |
| 7 | 28 | | | | | | | | 30 | 30 | 32 | 32 | | | | 32 | 32 | |
| 8 | 24 | | | 32 | 28 | 28 | | 32 | 32 | 32 | 32 | 34 | | | | 36 | 34 | |
| 9 | 24 | | | | | | | | 28 | 28 | 30 | 30 | | | 34 | 32 | 32 | |
| 10 | 26 | | 28 | 30 | 30 | 30 | | 30 | 32 | 32 | 36 | 34 | | | 36 | 34 | 36 | |

**B- Timed mating weight readings**

| Mice no. | D1 | D2 | D3 | D4 | D5 | D6 | D7 | D8 | D9 | D10 | D11 | D12 | D13 | D14 |
|---|---|---|---|---|---|---|---|---|---|---|---|---|---|---|
| 1 | 22 | | | 24 | 24 | | | 24 | 24 | 24 | 28 | 28 | | |
| 2 | 14 | | 14 | 14 | | | | 18 | 18 | 18 | 20 | 20 | | |

| | | | | | | | | | | | | | | |
|---|---|---|---|---|---|---|---|---|---|---|---|---|---|---|
| 3 | 26 | | 28 | 26 | 32 | | | 32 | 30 | 30 | 30 | 30 | | |
| 4 | 26 | 32 | 34 | 36 | 38 | | | 42 | 42 | 40 | 42 | 42 | | |
| 5 | 14 | | 14 | 14 | | | | 18 | 22 | 20 | 20 | 20 | | |
| 6 | 28 | | | 30 | 30 | | | 30 | 30 | 28 | 28 | 28 | | |
| 7 | 18 | | | | | | | | | | | | | |
| 8 | 30 | | | 30 | 30 | | | 36 | 38 | 38 | 38 | 40 | | |
| 9 | 24 | 26 | 26 | 28 | 29 | 32 | 32 | 30 | 30 | 32 | 34 | 32 | 32 | 32 |
| 10 | 22 | 24 | 24 | 26 | 27 | 28 | 28 | 30 | 32 | 36 | 36 | 38 | 40 | |

**C- Vaginal cytology and setting up of matings**

| Mice no | D1 | D2 | D3 | D4 | D5 | D8 |
|---|---|---|---|---|---|---|
| 1 | Diestrous | Diestrous | Diestrous | Estrous (Mating) | | |
| 2 | Metestrous | Metestrous | Metestrous | Metestrous | Diestrous | Estrous (Mating) |
| 3 | Diestrous | Diestrous | Metestrous | Diestrous | Diestrous | Estrous (Mating) |
| 4 | Estrous (Mating) | | | | | |
| 5 | Metestrous | Metestrous | Diestrous | Diestrous | Metestrous | Proestrous(Mating) |
| 6 | Diestrous | Diestrous | Diestrous | Proestrous (Mating) | | |
| 7 | Paralyzed | | | | | |
| 8 | Diestrous | Diestrous | Diestrous | Estrous(Mating) | | |
| 9 | Diestrous | Diestrous | Estrous ( mating) | | | |
| 10 | Diestrous | Metestrous | Proestrous (mating) | | | |
